# Source reconstruction of broadband EEG/MEG data using the frequency-adaptive broadband (FAB) beamformer

**DOI:** 10.1101/502690

**Authors:** Matthias Treder, Guido Nolte

## Abstract

A beamformer enhances the signal from a voxel of interest by minimising interference from all other locations represented in the sensor covariance matrix. However, the presence of narrowband oscillations in EEG/MEG implies that the spatial structure of the covariance matrix, and hence also the optimal beamformer, depends on the frequency. The frequency-adaptive broadband (FAB) beamformer introduced here exploits this fact in the Fourier domain by partitioning the covariance matrix into cross-spectra corresponding to different frequencies. For each frequency bin, an individual spatial filter is constructed. This assures optimal noise suppression across the frequency spectrum. After applying the spatial filters in the frequency domain, the broadband source signal is recovered using the inverse Fourier transform. MEG simulations using artificial data and real resting-state measurements were used to compare the FAB beamformer to the LCMV beamformer and MNE. The FAB beamformer significantly outperforms both methods in terms of the quality of the reconstructed time series. To our knowledge, the FAB beamformer is the first beamforming approach tailored for the analysis of broadband neuroimaging data. Due to its frequency-adaptive noise suppression, the reconstructed source time series is suited for further time-frequency or connectivity analysis in source space.

## Introduction

Electroencephalography (EEG) and magnetoencephalography (MEG), jointly abbreviated as MEEG here, measure the electrical currents or magnetic fields associated with synchronised activity of neuronal populations [1–3]. Compared to hemodynamic methods such as functional magnetic resonance imaging (fMRI), the Achilles’ heel of MEEG is the low accuracy in localising the sources. Since the sensors are located at a distance to the brain, the activity of brain sources needs to be statistically reconstructed. The inverse problem of estimating the sources from the sensor measurements is underconstrained because the number of sources exceeds the number of sensors. Furthermore, neurons in some subcortical areas are aligned such that they constitute a closed field, producing little measurable electrical activity [1]. Consequently, in order to obtain a unique solution, additional assumptions about the sources (such as their uncorrelatedness) are required to constrain the inverse model [4,5]. The linear forward model of MEEG, specifying how activity in the brain projects into the sensors, can be formulated as

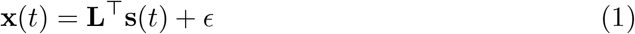

where **x**(*t*) ∈ ℝ^*N*^ is activity measured in *N* sensors at time *t*, **s**(*t*) ∈ ℝ^*M*^ is the source activity in *M* brain sources at time *t*, **L** ∈ ℝ^*M*×*N*^ is the *leadfield matrix* relating source activity to the EEG/MEG sensors, and *ϵ* ∈ ℝ^*N*^ is a noise term representing sensor noise and non-neural artefacts such as muscle activity. The i-th row of the leadfield matrix is called the *spatial pattern* of the i-th source. The leadfield matrix **L** is typically derived from forward calculations using a subsampled MRI or a cortical surface model [1]. Here, it is assumed that the dipole orientations are fixed, but the model is very similar for free dipole orientations (see below).

### Source localisation versus source reconstruction

Often, the primary purpose of source imaging is pinpointing the neural location that gives rise to observed EEG/MEG activity, i.e. *source localisation*. Dipole fitting models such as Multiple Signal Classification (MUSIC) and their extensions [6–8] search for a small number of sources that explain the data with sufficient accuracy.

Some imaging approaches such as minimum-norm estimates [9–11] and beamformers also provide linear functionals called *spatial filters* which allow for the estimation of the time series at a given voxel, i.e. *source reconstruction*. It has been shown that these methods can be cast into the generalised model

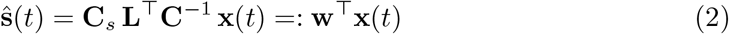

where **C** ∈ ℝ^*N*×*N*^ is the sensor covariance matrix, **C**_*s*_ ∈ ℝ^*M*×*M*^ is the source covariance matrix, 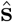 is the estimated source amplitude (a.k.a. virtual channel), and **w** ∈ ℝ^*N*^ is the spatial filter [12].

### Time-domain analysis: the LCMV beamformer

The purpose of beamforming is to optimally enhance the neuronal signal from a specific location in the brain with a known spatial pattern [3–5,13]. Several beamforming approaches have been shown to be equivalent to the *Linearly Constrained Minimum Variance (LCMV) beamformer* [14,15]. They differ mainly in the way that neural activity is quantified [13]. The spatial filter is given as

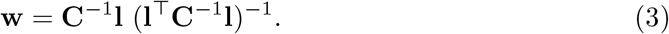

where *l* ∈ ℝ^*N*^ is the row of the leadfield matrix corresponding to the target source. The principal assumption for beamforming is that all sources are uncorrelated and hence the source covariance matrix **C**_*s*_ is diagonal. Indeed, inserting a diagonal source covariance matrix into Eq 2 leads to a scaled version of the beamformer solution in Eq 3. Recently, it has been shown that LCMV beamforming is formally equivalent to Linear Discriminant Analysis (LDA) [16].

For *source localisation*, source amplitude or variance can be calculated for all source locations. Peaks in the source power map indicate an active source. However, the relatively small norm of leadfields for deep sources causes a depth bias by amplifying noise. To account for this, the source power estimate is usually divided by noise power. The resulting quantity was originally introduced as *neural activity index* by Van Veen et al. [15] and is given here for fixed dipoles [13,17] as

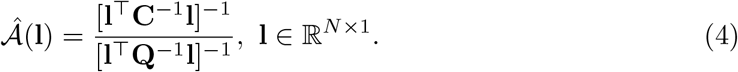

For free dipoles, the spatial pattern **l** = [**l**_*x*_ **l**_*y*_ **l**_*z*_] consists of 3 components along the x, y, and z axes. In the original approach by Veen et al., unreliable directions with a small norm can dominate the resulting quantity. The modified index by Huang et al. [13] remedies the problem by normalising each direction separately:

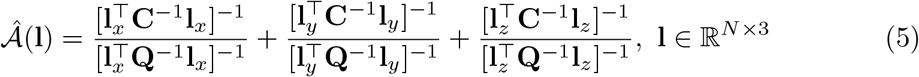

The noise covariance matrix **Q** can be estimated e.g. by using pre-stimulus activity in an event-related design or empty room recordings in the case of MEG.

### Frequency-domain analysis: DICS

Beamforming in the frequency domain is implemented by the *Dynamic Imaging of Coherent Sources (DICS)* approach [18,19]. The cross-spectral density matrix **C**(*ω*) is determined for a particular frequency *ω*. The real part of **C**(*ω*) then takes the role of the covariance matrix, leading to a frequency-dependent solution analogous to LCMV beamforming:

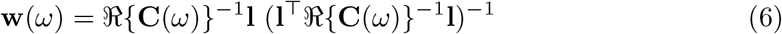

where ℜ(·) designates the real part. Beamforming in the time-frequency domain was introduced by Dalal et al. [20]. Sensor time series are passed through filter banks enabling frequency-specific beamformers. Additionally, cross-spectra are calculated for specific time segments, accounting for nonstationarity. As a result, beamformer weights are obtained for each time-frequency bin

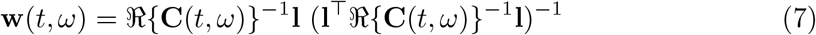

where the spatial filter and the cross-spectral density are now functions of both time *t* and frequency *ω*.

### Towards broadband beamforming

What is the optimal spatial filter for reconstructing a broadband signal? As illustrated in Fig 1, the spatial structure of the background activity varies as a function of frequency. Clearly, a spatial filter that is optimal for a given frequency band is most likely not optimal for a different frequency band.

**Fig 1.**
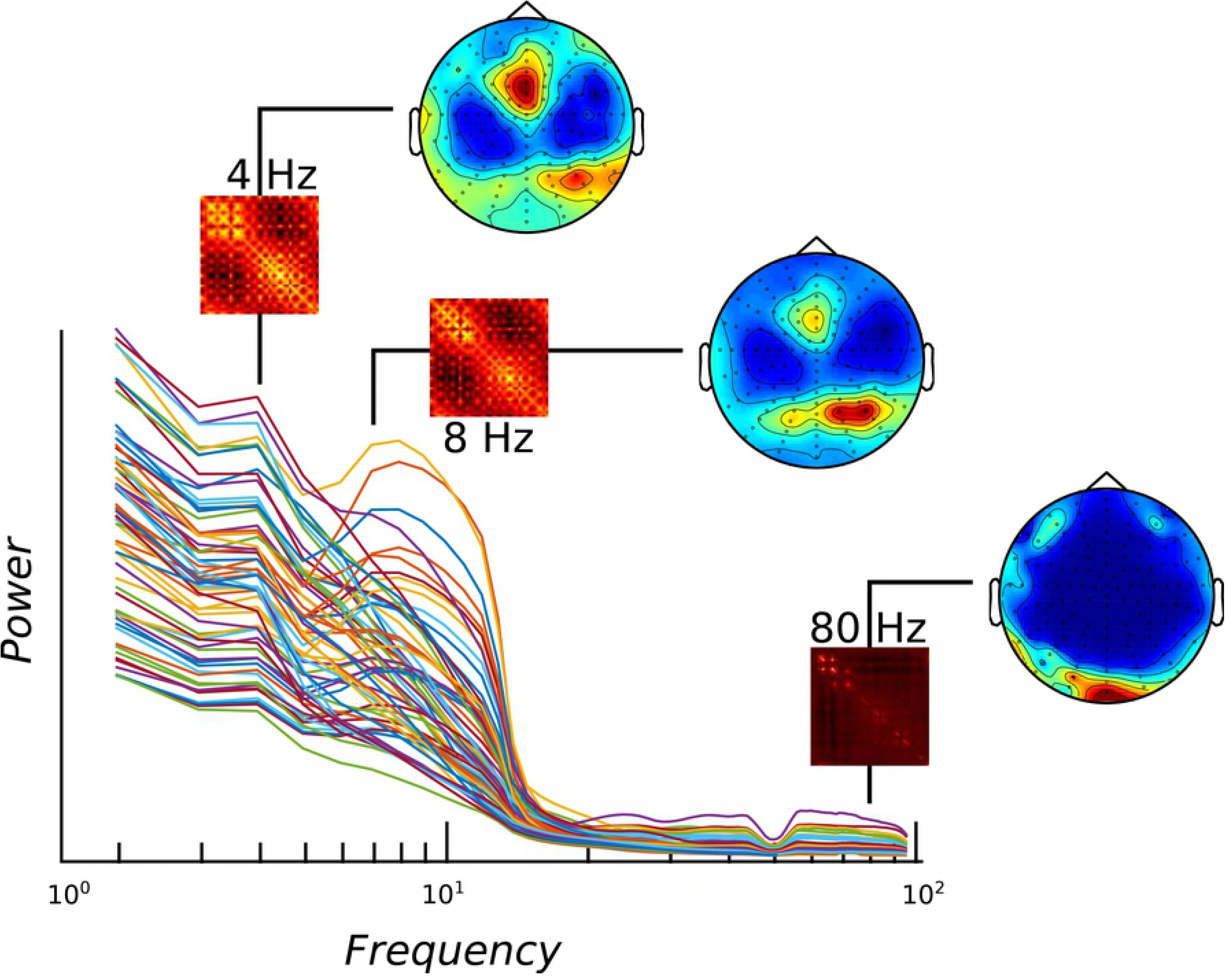
Power spectra. Power spectra for different electrodes (coloured lines) for an example EEG dataset, depicted on a logarithmic frequency axis. For the frequencies 4, 8, and 80 Hz, the real parts of the cross-spectra are shown as small rectangles. To better illustrate the spatial structure of the cross-spectra, each topography depicts the eigenvector of the covariance matrix corresponding to the largest eigenvalue with magnitude being colour-coded. Clearly, the spatial structure of the cross-spectrum changes as a function of frequency.

The LCMV beamformer, when applied to broadband data, is biased towards cancelling noise in low frequencies because they contribute most of the variance [20], producing a flattened frequency spectrum of the estimated source [21]. DICS, on the other hand, has an excellent signal-to-noise ratio because beamforming is restricted to a narrow frequency band. The disadvantage is that DICS cannot be directly applied to broadband signals. The time-frequency beamformer solves this problem by splitting the signal into time-frequency bins [20]. However, it is computationally expensive and the different frequencies are analysed separately.

Here, a different approach is taken by considering a broadband signal in the frequency domain. Due to the linearity of the Fourier transform, applying a spatial filter in the time domain is equivalent to applying it in the Fourier domain. However, instead of using the same spatial filter for each frequency, one can divide the spectrum up into frequency bins and calculate the cross-spectrum corresponding to each bin. This allows for the construction of a different spatial filter for each frequency bin. After applying these spatial filters, the Fourier transform can then be inverted to recover the broadband signal in source space. We call this approach *frequency-adaptive broadband (FAB) beamforming*.

At this point, the investigator can proceed with any time or frequency domain analysis, such as event-related potentials, single-trial decoding, time-frequency decomposition, or connectivity analysis. Furthermore, by combining power estimates across frequency bins, FAB beamforming can also be used for source localisation of broadband data.

## Materials and methods

### Partitioning of the broadband covariance matrix

A convenient property of the Fourier transform is that it partitions the time-domain covariance matrix into frequency-specific cross-spectra. The existence of narrowband oscillations in MEEG assures that this partitioning is non-trivial, that is, the cross-spectra have different spatial structures (see Fig 1). Analogous to minimising the covariance matrix by a single spatial filter (as in LCMV beamforming), one can instead perform beamforming by minimising the cross-spectra.

Let **X** ∈ ℝ^*N*×*T*^ be the sensors × time data matrix, and let **x**_*i*_ ∈ ℝ^*T*^ be the i-th row of **X** corresponding to the i-th MEEG channel. Then its Fourier transform is defined as 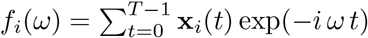 and *x_i_*(*t*) can be expressed using the inverse Fourier transform as 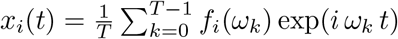 where the frequency index *k* is related to the angular frequency as *ω_k_* = 2*πk*/*T* for *k* = 0, 1, …, *T* – 1.

Assuming that all channels have zero mean, the covariance between the i-th and the j-th channel, corresponding to the (ij)-th entry of the covariance matrix **C**, and their cross-spectral density, corresponding to the (i,j)-th entry of the cross-spectral density matrix **C**(*ω*) for frequency *ω* can be defined as

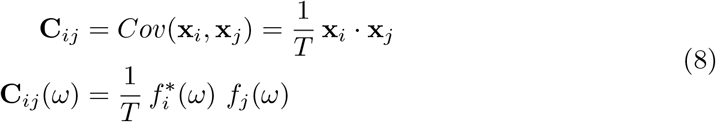

where · denotes the complex dot product. Using the orthogonality of the Discrete Fourier transform, 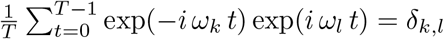, and inserting the inverse Fourier transform into Eq 8 one can express the dot product of **x**_*i*_ and **x**_*j*_ by the dot product of their Fourier coefficients as

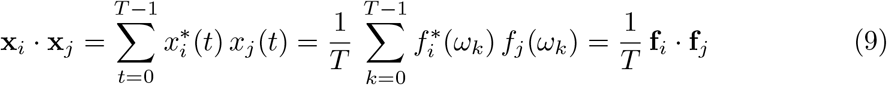

Together, Eq 8 and Eq 9 show that the cross-terms vanish and hence the broadband covariance matrix can be partitioned into cross-spectral densities:

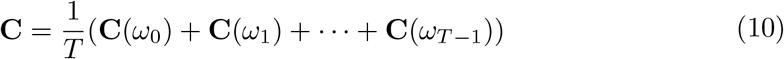

Now, the conjugate symmetry of the Fourier spectrum of real signals can be exploited, implying that **C**(*ω*_*k*_) = **C**(*ω*_*T*–*k*_)*. Hence, the covariance matrix can be expressed by the real parts of the cross-spectra as

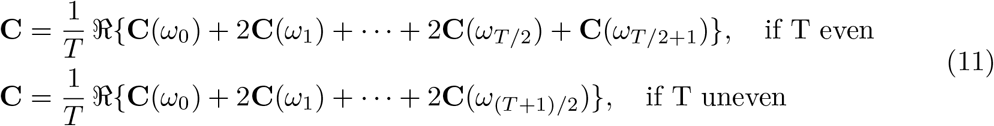

### Estimation of the cross-spectrum

The periodogram is an unbiased but inconsistent estimator of the power spectral density. In Welch’s method, the signal is divided up into overlapping segments. Each segment is multiplied by a window function and then DFTs are calculated. The cross-spectra derived from the individual segments are then averaged. This significantly improves the estimate at the expense of frequency resolution. However, a resolution of e.g. less than 0.5 Hz is rarely needed for neural oscillations in MEEG, justifying this trade-off. An alternative way of spectrum estimation is the use of multitapers. Here, a Slepian sequence of mutually orthogonal tapers is used to obtain independent power estimates [22].

### Regularisation

EEG and MEG signals are low-dimensional, often leading to ill-conditioned covariance and cross-spectral density matrices. Moreover, if a subspace has been removed in the course of artefact correction (e.g. using maxfiltering or Independent Components Analysis), the cross-spectral matrices are singular. To guarantee invertibility, they can be slightly manipulated using a regularisation approach. In analogy to shrinkage estimation in covariance matrices [23–25], the shrinkage regularisation approach is adapted here to the cross-spectra:

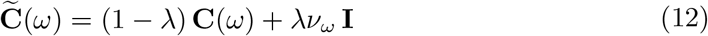

Here, **I** ∈ ℝ^*N*×*N*^ is the identity matrix and *ν_ω_* = trace(**C**(*ω*))/*N* scales the identity matrix such that its trace matches the trace of **C**(*ω*). *λ* is the regularisation parameter that blends between the empirical cross-spectrum (*λ* = 0) and a diagonal matrix with equal total variance (*λ* = 1). Typically, a small value of *λ* in the range of 10^−3^ is sufficient to ensure invertibility without distorting the empirical cross-spectrum too much. Note that regularisation does not need to come at the expense of a bad estimate. In some cases, the bias introduced by regularisation can even increase the accuracy of the inverse model [24,26].

### Frequency-adaptive broadband (FAB) beamforming

Using the regularised cross-spectra, the frequency-adaptive spatial filter for frequency *ω* is given by

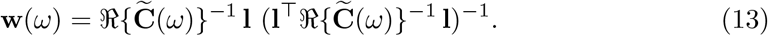

Let **F** ∈ ℝ^*N*×*T*^ be the matrix containing the [1 × *T*] Fast Fourier Transforms (FFTs) of the sensor time series as rows, and let **F**(*ω*) ∈ ℝ^*N*^ be the column of **F** corresponding to the frequency *ω*. The source-space Fourier coefficient for frequency *ω* is then given by

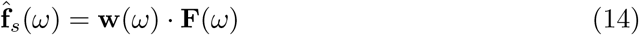

Note that in practice the cross-spectra are calculated across multiple segments, leading to a lower frequency resolution but better estimate. For the purpose of obtaining the Fourier coefficients of the source, the spatial filter **w**(*ω*) can therefore be applied in the frequency neighbourhood [*ω* – *ϵ*/2, *ω* + *ϵ*/2], where *ϵ* is the width of the frequency bins. Once Fourier coefficients for all frequencies have been calculated, the inverse Fourier transform is calculated to recover the source time series.

In order to use the FAB beamformer for *source localisation*, information needs to be accumulated across the different frequency bins. In Eq 4, the neural activity index was introduced as a way to represent signal power normalised by noise power. As a straightforward extension of this idea, the *summed neural activity index* can be introduced, which essentially sums the neural activity scores across the frequency bins to measure the collective evidence for the presence of a signal. Here, the formula is given for fixed dipoles, but the extension to free dipoles is straightforward:

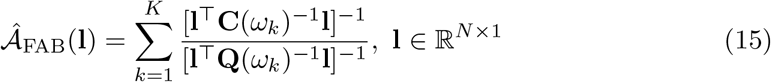

It is assumed that the signal is analysed in *K* different frequency bins, and **Q**(*ω*_*k*_) is the noise cross-spectrum at frequency *ω*_*k*_. Due to the 1/f shape of the MEEG power spectrum, it can be expedient to use a logarithmic spacing of the frequency bins. This is because with a linear spacing, higher frequency bands such as the gamma band would dominate the outcome by contributing more frequency bins than the lower frequency bands.

### Simulations

To evaluate the efficacy of the FAB beamformer in reconstructing sources, simulations were performed wherein it was compared to LCMV beamforming and minimum-norm estimates (MNE) [9,10,27,28]. Simulations were based on artificial signals in source space (simulation 1) or real MEG data from a resting-state measurement (simulation 2). To build source models, individual MRI scans of 582 subjects that took part in the Cambridge Center for Ageing and Neuroscience (Cam-CAN) study on healthy ageing [29–31] were used. See [29] for a detailed protocol of the study and [30] for the processing pipeline.

#### Data acquisition and preprocessing

MRI data were acquired at the MRC Cognition and Brain Sciences Unit (MRC-CBU), Cambridge, UK, using a Siemens 3T TIM TRIO (Siemens, Erlangen, Germany) with a 32-channel head coil. Anatomical images were measured with a resolution of 1mm^3^ isotropic using a T1-weighted MPRAGE sequence (TR: 2250ms; TE: 2.98ms; TI: 900ms; 190Hz; flip angle: 9°; FOV: 256 × 240 × 192mm; GRAPPA acceleration factor: 2; acquisition time of 4 minutes and 32 seconds), and a 1mm^3^ isotropic T2-weighted SPACE sequence (TR: 2800ms; TE: 408ms; flip angle: 9°; FOV: 256 × 256 × 192mm; GRAPPA acceleration factor: 2; acquisition time of 4 minutes and 32 seconds).

MEG data were collected continuously using a whole-head Elekta Neuromag Vector View 306 channel MEG system with 102 magnetometers and 204 planar gradiometers (Elekta, Neuromag, Helsinki, Finland), located in a light magnetically shielded room at the MRC-CBU. Data were sampled at 1kHz and high-pass filtered at 0.03Hz. Head motion was corrected offline based on 4 Head-Position Indicator (HPI) coils. Approximately 8 minutes and 40 seconds of eyes-closed resting state data was acquired from each participant, with the first 20 seconds discarded from further analysis. Ethical approval for the study was obtained from the Cambridgeshire Research ethics committee. A 3D digitizer (Fastrak Polhemus, Colchester, VA) was used to measure the positions of the HPI coils, approximately 100 head-shape points across the scalp, and three anatomical fiducials (nasion, left and right preauricular points).

MEG data was then preprocessed using MaxFilter 2.2.(Elekta Neuromag Oy, Helsinki, Finland). It uses spherical basis functions to reject environmental noise based on magnetic fields outside a sphere enclosing the sensor array, by removing temporally correlated sources inside this sphere but outside a sphere enclosing the brain using the temporal extension of signal space separation (tSSS) [32], with a correlation threshold of 0.98 and a 10 s sliding window. The tSSS was further used to perform movement correction every 200 ms using the HPI coils data, and to align all participants’ data in a common space (as if the centre of their heads were in the same position relative to the sensors). Finally, MaxFilter was also used to detect and reconstruct bad channels, and to notch-filter the line noise at 50 Hz and its harmonics. ICA was then used to identify physiological artefacts from blinks, eye-movements, and cardiac pulse. This was done by identifying those components whose time courses and spatial topographies correlated highly with reference time courses (correlation greater than three standard deviations from mean) and spatial topographies (correlation greater than two standard deviations from mean), respectively, for each of the above artefact types.

#### Source modelling

Structural MRI images were processed using automated segmentation algorithms in FreeSurfer (http://surfer.nmr.mgh.harvard.edu/) [33,34]. The white-matter surface was extracted for further source modelling. The MNE software was used with the ico-2 spacing option in order to decimate the white-matter surface to 324 vertices (162 per hemisphere) [35]. Surfaces were coregistered with the MEG sensors using FieldTrip [36]. An initial alignment was performed using the fiducial markers (nasion, left and right preauricular points). The coregistration was then refined using the headshape points. Forward modelling for 204 planar gradiometers was performed using a realistic single-shell volume conductor [37]. This yielded a set of spatial patterns in x, y, and z directions for each vertex collected in a *N* × 3 matrix. To obtain sources with fixed dipoles, a singular value decomposition (SVD) was performed on the matrix for each vertex. The principal component corresponding to the maximum eigenvalue was then chosen as fixed dipole. There was a total of 582 subjects, aged 18-88, for which source spaces had been created in this way.

#### Inverse modelling

For the LCMV beamformer, Eq 3 was used along with a regularised covariance matrix using shrinkage. For MNE, the regularised inverse operator can be defined as

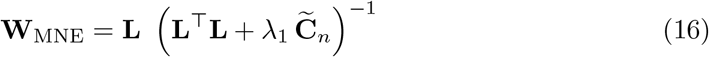

where 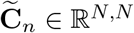 is the regularised noise covariance matrix and *λ*_1_ is a regularisation parameter. For the simulations, the noise covariance matrix **C**_*n*_ was extracted from empty room recordings and then regularised as 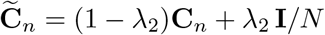, where *λ*_2_ is another regularisation parameter and **I** is the identity matrix. To bring the matrices on equal footing, **L**^⟙^**L** and **C**_*n*_ were both scaled to have a trace of 1. Furthermore, for convenience, a convex regularisation term 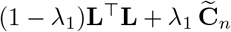 was used in Eq 16 for the simulations. Except for scaling, it does not affect the solution, but it allows to define the regularisation parameter *λ*_1_ on the closed interval [0,1], just like the other regularisation parameters.

#### Simulation 1: Reconstruction of simulated MEG

Six minutes of MEG data sampled at 256 Hz were simulated at each of the 324 vertices in source space. First, one of the vertices was randomly selected as “signal vertex” representing the signal of interest. The other vertices were used as noise. Noise source were created as band-limited signals in the Fourier domain: First, the power spectrum of the noise source was modelled by a Hanning window that had a random location in frequency space and a random band-width of up to 64 Hz. Each Fourier coefficient was given a random phase and the resulting spectrum was then inverse Fourier-transformed into the time domain. As target signal, either a band-limited oscillation created the same way as the noise was used or alternatively a 1/f (‘pink’) signal. Using the forward models, signal and noise sources were mixed together into the 204 planar gradiometers. This is illustrated in Fig 2.

**Fig 2.**
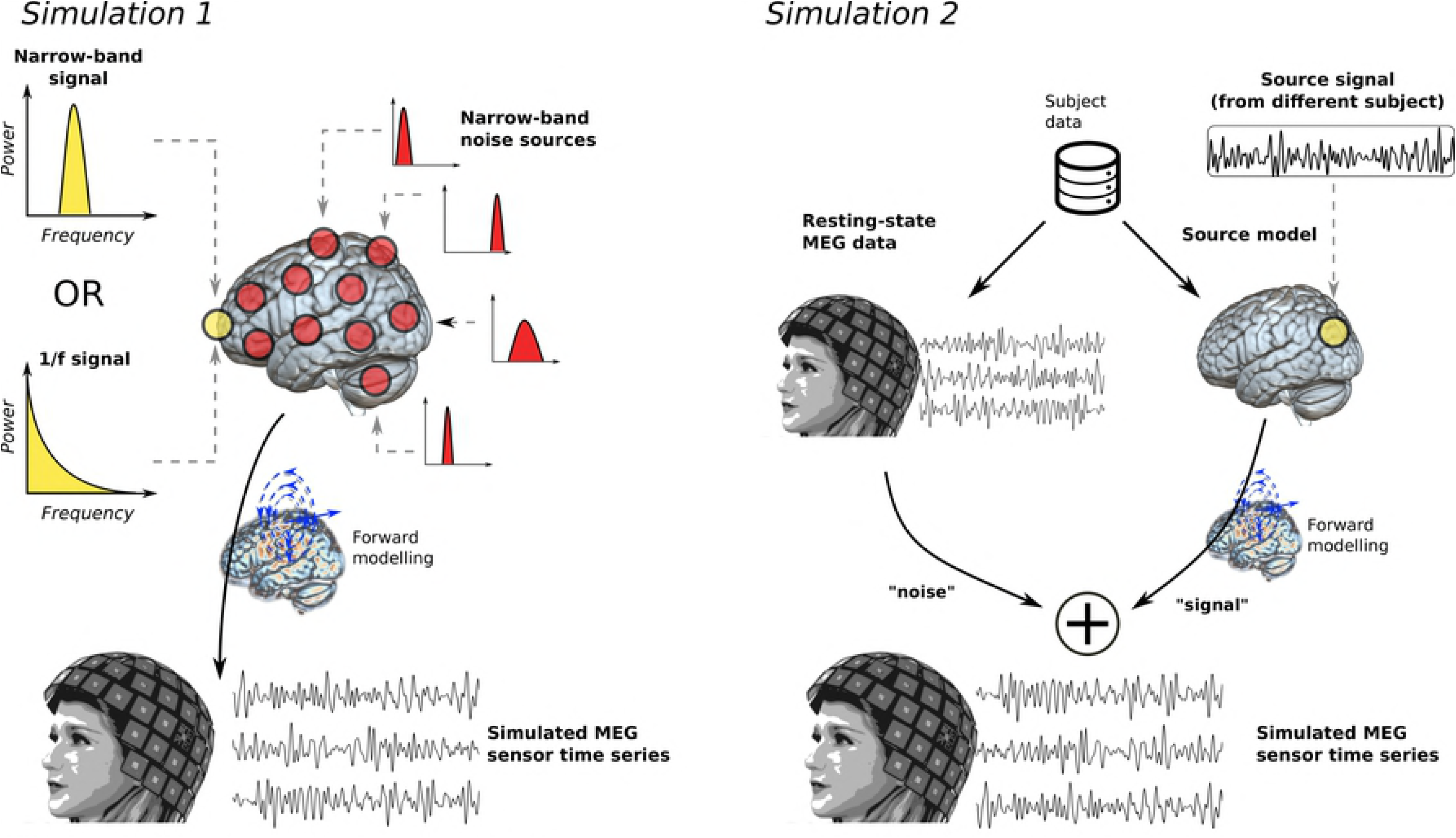
Simulation data. Graphical illustration of how the data for the simulations was created. In simulation 1, real source spaces were used together with artificially generated signals consisting of a narrowband or 1/f signal (yellow power spectra) and narrowband noise sources (red). The signals were mapped to sensor space using forward models. In simulation 2, real resting-state measurements were used as background noise. In order to have a ground truth, additionally a source time series from another subject was used as the target signal. It was mapped to sensor space using the first subject’s source space and then added to the resting-state data.

Separate simulations were performed for every of the 582 subjects, using the source spaces based on their individual MRIs. Furthermore, simulations were repeated 20 times for every subject, with a randomly chosen signal vertex and randomly generated signals in each iteration, yielding a total of 582 · 20 = 14, 550 MEG simulations. Signal-to-noise ratio (SNR), defined in sensor space as the trace of the signal covariance divided by the trace of the noise covariance, was increased from 10^−6^ to 10^0^ in 25 logarithmic steps.

Finally, to ensure that any difference between the inverse models is not due to a specific choice of the regularisation parameters (the *λ*’s), grid searches were conducted for optimising the parameters. Ten different values for lambda were tested in logarithmic steps, i.e. *λ* = 10^−9^, 10^−8^, …, 10^0^. For the FAB beamformer, *λ* determined the regularisation of the cross-spectral matrices. For the LCMV beamformer, *λ* determined the regularisation of the covariance matrix. For the MNE, a two-dimensional search grid for *λ*_1_, *λ*_2_ as introduced above was used.

In each iteration, and for each SNR and *λ* value, the time series in the signal vertex was reconstructed using the three inverse models. For the FAB beamformer, cross-spectra were calculated by splitting the MEG into 1s segments with 50% overlap. Each segment was multiplied by a Hanning window and an FFT was performed, yielding a spectral resolution of 1 Hz. The cross-spectra were then averaged across segments. This way, cross-spectra were obtained for all frequencies between 1 and 100 Hz in 1 Hz steps. For the LCMV beamformer, the covariance matrix was calculated from the broadband signal. To measure reconstruction quality, we used Pearson correlation coefficients between the reconstructed time series and the known true time series.

#### Simulation 2: Resting-state MEG

In simulation 1, real source models and artificially generated time series with narrowband spectra were used in order to illustrate the reconstruction quality of the FAB beamformer. However, these results do not necessarily generalise to true MEEG signals, due to the idiosyncratic properties of MEEG such as the 1/f power spectrum.

To investigate whether the FAB beamformer outperforms the other methods in real MEG data, resting-state data downsampled to 200 Hz was used as described above. In order to be able to quantify reconstruction quality, a ‘ground truth’, i.e. a known source time series was required. To this end, a source time series obtained with MNE from a second, randomly selected subject and a randomly selected vertex was used. This time series was assigned to a randomly chosen vertex and then forward projected into the sensor space of the first subject using the first subject’s forward model. This is illustrated in Fig 2.

To reduce computational load, simulations were performed for a medium SNR value of 10^−3^ for which FAB, LCMV and MNE differed well in simulation 1. SNR was defined as the trace of the multivariate signal (source signal of second subject mapped to sensor space) divided by the trace of the resting-state data. Since SNR was defined on basis of the broadband data and the power varies substantially as a function of frequency, the effective SNR was somewhat different for each frequency band. However, this was not a problem since we contrasted the different methods within the frequency bands.

For FAB and LCMV, the regularisation parameter was fixed to *λ* = 10^−3^ because it was the largest value that had been selected in simulation 1. For MNE, which performed worse than the other two methods in simulation 1, a grid search was again performed in order to maximise MNE performance. Grid search is less costly for MNE since the spatial filters are independent of the MEG data and thus have to be computed only once for every subject. For every subject, the simulation was repeated 25 times. In each iteration, a different source time series from a randomly selected second subject was used as signal.

Additionally, to investigate in how far the performance of the FAB beamformer depends on the cross-spectral estimation method, we compared FFT-based estimation using Hanning windows (used in simulation 1) to spectral estimation based on multitapers and Morlet wavelets. To this end, the MEG was split into 2s segments with 50% overlap. Multitaper-based estimation was performed using discrete prolate spheroidal sequences (Slepian sequences) with 5 tapers. Cross-spectra were obtained for all frequencies between 1 and 100 Hz in 1 Hz steps and then averaged across tapers and across segments. Wavelet-based estimation was performed using Morlet wavelets in the Fourier domain. The FFT of each segment was tapered with a Gaussian window, the Fourier domain representation of a Morlet wavelet. The time-frequency trade-off was determined by setting wavelet width to 5. For instance, at a frequency of 10 Hz, this trade-off leads to a bandwidth of 10/5*2 = 4 Hz. The cross-spectrum was then calculated by summing across all non-zero frequency bins. Cross-spectra were obtained for all frequencies between 1 and 100 Hz in logarithmic steps and then averaged across segments.

The resulting broadband source signals were bandpass filtered in the delta (1–4 Hz), theta (4–8 Hz), alpha (8–12 Hz), beta (12–30 Hz), lower gamma (30–60 Hz), and upper gamma (60–99 Hz) band. Reconstruction quality was again quantified using Pearson correlation between the bandpass filtered true source time series and the bandpass filtered reconstructed time series, separately for each frequency band, each inverse method and each of the three methods for calculating the cross-spectrum. Finally, we also compared the FAB beamformer to frequency-specific LCMV beamformers that were applied to bandpass-filtered data.

## Results

### Simulation 1

For every subject, all correlation coefficients were first Fisher-z transformed. Subsequently, for each SNR and *λ* value, medians of the z-values were calculated across the iterations. The optimal regularisation parameters were determined by inspecting the maximum z-values summed across all SNR levels. The z-values corresponding to the optimal parameters were transformed back to correlation values using the hyperbolic tangent function, obtaining a correlation coefficient for each SNR level.

Fig 3 shows the results of simulation 1. The FAB beamformer yields a better reconstruction quality than either of the other two methods for most SNR levels. The inverse methods converge only for very high or low levels of SNR. To test statistical significance, two different 3-way analyses of variance (ANOVA) were performed on the Fisher-z transformed correlation values, one for each of the competing methods (LCMV and MNE). *α* = 0.025 was used to correct for two comparisons. Inverse method, signal type (narrowband, 1/f), and SNR level served as factors. For the ANOVA comparing FAB and LCMV beamforming, SNR (*F* = 260.41, *p* < .0001) and inverse method (*F* = 43.07, *p* < .0001) yielded significant results, but signal type (*p* = 0.08) did not. In the second ANOVA comparing the FAB beamformer to MNE, SNR (*F* = 241.13, *p* < .0001) and inverse method (*F* = 47.89, *p* < .0001) yielded significant results, but signal type (*p* = 0.1) did not. Hence, the FAB beamformer globally outperformed both the LCMV beamformer and MNE in terms of reconstruction quality.

**Fig 3.**
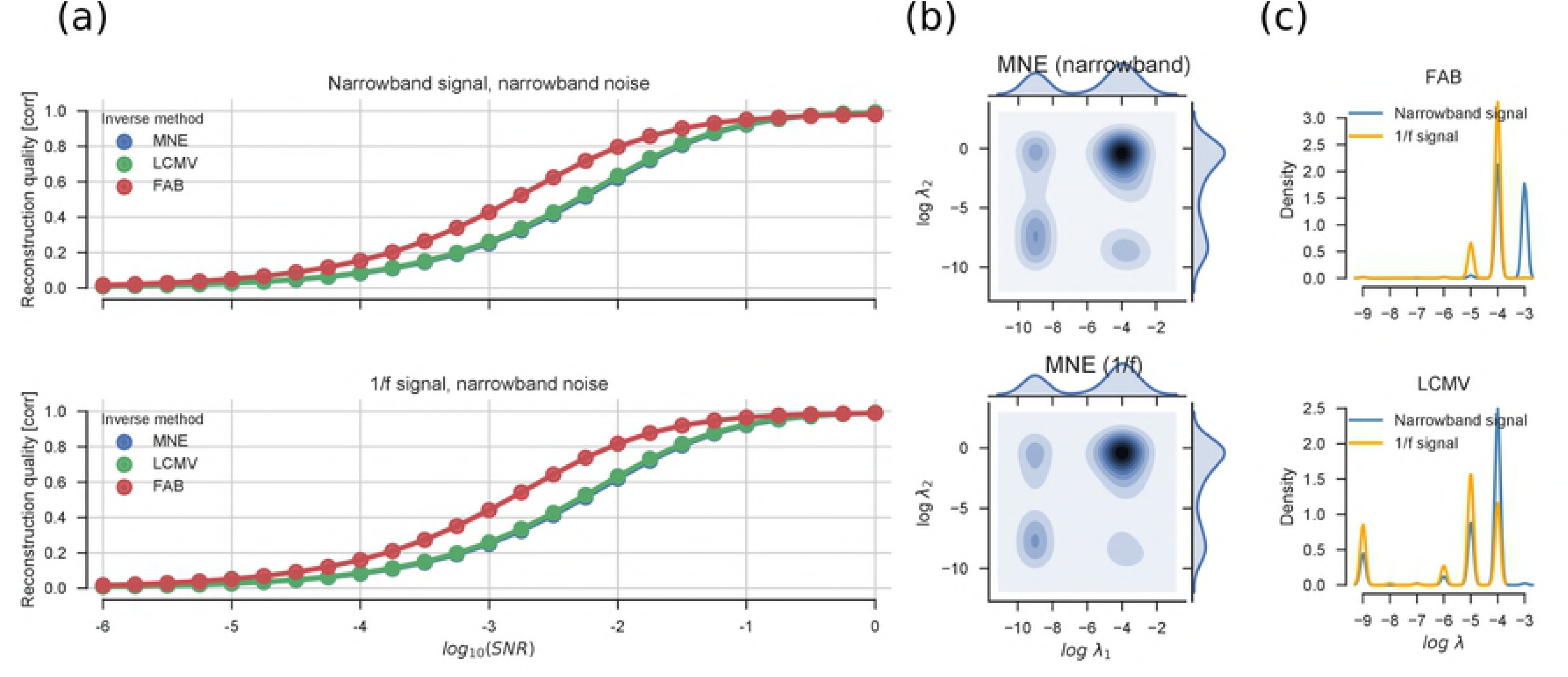
Results of simulation 1. (a) Reconstruction quality, measured as correlation between true source signal and the source reconstructed signal a for narrowband signal (top) and a 1/f signal (bottom). The standard error is too small to be displayed. Note that the results for LCMV beamformer (green) and MNE (blue) are almost identical, which is why the MNE data is not easily seen. (b) Bivariate distribution of best regularisation parameters *λ*_1_ and *λ*_2_ across subjects for MNE. (c) Distributions of best regularisation parameter *λ* for FAB and LCMV beamformer.

The regularisation parameters have different meanings for different methods, so their distributions were not subjected to hypothesis testing. However, one can qualitatively appreciate that for MNE, the bivariate distribution of *λ*_1_ and *λ*_2_ shows multiple modes. Mostly, large regularisation values were selected for both parameters. In general, the *λ*’s tend to be either large or small, with a low occurrence of intermediate values. The parameters for LCMV and FAB beamforming are comparable to some extent since they both refer to the data covariance. For both beamformers, regularisation tends to be stronger for narrowband signals (blue line) than for 1/f signals (orange). At least for the FAB beamformer, this may be partially explained by the fact that 1/f signals contribute variance at all frequencies, whereas for narrowband signals the information is concentrated at a specific frequency, requiring more regularisation for the other frequency bins. Furthermore, the strength of regularisation in LCMV beamforming tends to be an order of magnitude smaller than in FAB beamformer. This could be due to the fact that the LCMV covariance matrix might be slightly better conditioned because it combines information across all frequency bands.

### Simulation 2

Fig 4 shows the results of simulation 2. Like in simulation 1, the correlation coefficients were preprocessed and averaged using Fisher-z transformations.

**Fig 4.**
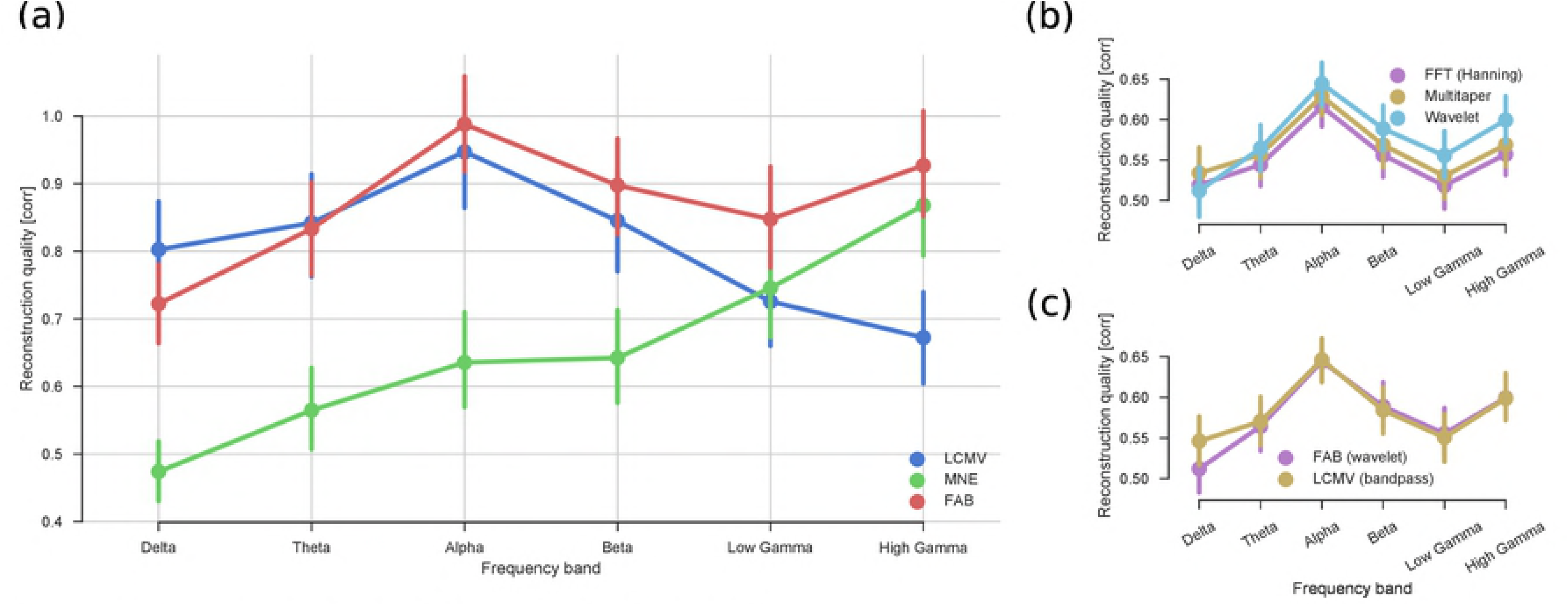
Results of simulation 2. Results of simulation 2 using resting-state MEG data. (a) Reconstruction quality of the broadband signal for different frequency bands. In medium to high frequencies, the FAB beamformer with wavelet-based estimation outperforms both the LCMV beamformer and MNE. (b) Comparing different ways of calculating cross-spectra in the FAB beamformer. The wavelet-based method outperforms multitapers, although it underperforms for low frequencies. Multitapers in turn outperform estimation based on FFT. (c) Comparing the FAB beamformer using wavelet-based estimation to separate LCMV beamformers that have been applied to bandpass-filtered data. Except for the theta band, where FAB underperforms, a single FAB beamformer is as good as separate LCMV beamformers.

It was then tested whether the FAB beamformer performs better than the other two methods in terms of reconstruction quality across different frequency bands (Fig 4a). To test statistical significance, two different 2-way analyses of variance (ANOVA) were performed on the Fisher-z transformed correlation values, one for each of the two competing methods (LCMV and MNE). Inverse method, frequency band and their interaction served as factors. The wavelet-based implementation of the FAB beamformer was used. For the ANOVA comparing FAB and LCMV beamforming, effects of frequency band (*F* = 7.84, *p* < .0001) and inverse method (*F* = 8.53, *p* < .01) were significant, as well as their interaction (*F* = 4.7, *p* < .001). The second ANOVA comparing the FAB beamformer to MNE yielded significant effects of frequency band (*F* = 17.01, *p* < .0001) and inverse method (*F* = 110.79, *p* < .0001) as well as a significant interaction (*F* = 4.95, *p* < .001).

To investigate whether the exact cross-spectral estimation procedure significantly affects FAB beamformer performance, a 2-way ANOVA with factors frequency band and estimation method (FFT, multitaper, wavelet) was performed (Fig 4b). There were significant effects of frequency band (*F* = 23.99, *p* < .0001) and estimation method (*F* = 12.47, *p* < .0001). Further ANOVAs contrasting pairs of estimation method showed that wavelet estimation is significantly better than multitapers (*F* = 16.91, *p* < .0001), and multitaper estimation is significantly better than FFT (*F* = 7.47, *p* < .0001).

Finally, we compared the wavelet-based FAB beamformer to LCMV beamformers applied to the bandpass-filtered data instead of the broadband data using a 2-way ANOVA. The FAB beamformer tended to be worse, but this was not significant (p = .06). The effect was mostly caused by differences in the theta band (Fig 4c), whereas FAB beamformer and bandpass LCMV beamformer were nearly on par for the other frequency bands.

## Discussion

A frequency-adaptive broadband (FAB) beamforming approach was introduced for the source reconstruction of broadband MEEG data. The inverse model was benchmarked against two state of the art competitors, namely the LCMV beamformer and MNE. The Pearson correlation between the known true time series and the reconstructed source was used as reconstruction quality measure.

In simulation 1, using narrowband noise sources and a single narrowband or 1/f target signal, the FAB beamformer consistently outperformed both the LCMV beamformer and MNE for a large range of SNR values (Fig 3). The methods converged only for very high or very low SNR values. At low SNR values, reconstruction quality in terms of Fisher-z values increased by up to 90% from FAB as compared to MNE/LCMV.

In simulation 2, each subject’s resting-state MEG served as real background activity. A source time series from a different subject was used as the target signal. Applied to broadband data, the LCMV beamformer reconstructed lower frequency bands very well but failed to efficiently reconstruct gamma activity (Fig 4). This is not surprising, since the broadband covariance matrix is dominated by lower frequencies. Vice versa, MNE performed relatively poor for low frequency bands but outperformed the LCMV beamformer in the gamma band. Crucially, the FAB beamformer showed a better overall performance than the other two methods, although it performed on par with the LCMV beamformer in the theta band and tended to be worse in the delta band.

Furthermore, the effect of spectral estimation procedure on FAB beamformer performance was investigated (Fig 4b). In terms of performance, it was found that wavelet > multitaper > FFT, although the wavelet-based estimation was underperforming for the delta band. This is possibly due to the logarithmic scaling of the frequency bandwidth of Morlet wavelets that does not match the true bandwidth of neural oscillations. Smoothing across frequency bins for low frequencies could increase frequency bandwidth and thereby alleviate this problem. Alternatively, the time-frequency trade-off could be changed towards a more coarse frequency resolution. Apart from this, the overall superior performance of the wavelet-based approach suggests that the distribution of information is broader at higher frequencies.

Crucially, except for the delta band, the FAB beamformer performed equally well as separate LCMV beamformers tested on the standard frequency bands (Fig 4c). This implies that the broadband time series obtained with the FAB beamformer has a high signal-to-noise ratio at every frequency.

## Conclusion

The FAB beamformer reconstructs the broadband time series of a vertex that has a high SNR across the frequency spectrum, unlike DICS and LCMV which are optimal only for a frequency band. Therefore, FAB beamforming can serve as a first step for further analysis in source space, for instance decoding of broadband ERPs or connectivity analysis. It might also be useful for representing neural events with complex neural signatures. For instance, during sleep K-complexes are often followed by sleep spindles. A single frequency band might not be able to capture the full dynamics of such a complex signal. Finally, in this paper we focused on source reconstruction. Further research is required to investigate the usefulness of the FAB beamformer for source localisation of broadband signals.

## Acknowledgements

This research was partially funded by the BMBF (031A130), the German Research Foundation (DFG, SFB936/A3/Z3 and TRR169/B1/B4), and from the Landesforschungsförderung Hamburg (CROSS, FV25).

